# The structural context of mutations in proteins predicts their effect on antibiotic resistance

**DOI:** 10.1101/2025.09.23.676583

**Authors:** Anna G. Green, Mahbuba Tasmin, Roger Vargas, Maha R. Farhat

**Affiliations:** Department of Biomedical Informatics, Harvard Medical School, 25 Shattuck St., Boston, MA, 02115, USA; Manning College of Information and Computer Sciences, University of Massachusetts, 140 Governors Dr., Amherst, MA, USA; Division of Pulmonary & Critical Care, Massachusetts General Hospital, 55 Fruit St, Boston, MA, 02114, USA

## Abstract

In *Mycobacterium tuberculosis,* a prevalent and deadly pathogen, resistance to antibiotics evolves primarily through non-synonymous mutations in proteins. Sequence-based analyses are currently used to understand the genetic basis of antibiotic resistance, either via genotype-phenotype association, or via signals of convergent evolution. These methods focus on primary sequence and often neglect other biological signals such as protein structural information. We hypothesize that integrating the structural context of mutations improves the prediction of effects on function and phenotype. We curate high confidence structural annotations for the *M. tuberculosis* proteome from 1,371 crystallography and 2,316 AlphaFold predictions, and combine the structures with mutations from over 31,000 clinical *M. tuberculosis* isolates. We demonstrate that mutations in proteins known to cause resistance are clustered in 3D space, even in proteins where inactivating mutations at any position are thought to cause resistance. We develop a statistic to search the *M. tuberculosis* proteome for signal of clustered mutations, finding over 450 proteins that display this signal, many of which have a known relationship with antibiotic resistance. We show that a supervised classifier trained on 3D distance to known resistance sites alone has an F1 score of 94.6% at classifying mutations as resistance-conferring across proteins. This work demonstrates that protein structure provides useful information for categorizing which variants may cause antibiotic resistance, even when the majority of structures are AI-predicted.

## Introduction

The increasing prevalence of antibiotic-resistant *Mycobacterium tuberculosis* is challenging control of tuberculosis (TB), a disease responsible for the highest number of infectious disease related deaths globally.^1^ Currently, diagnosis of antibiotic-resistant tuberculosis relies on time-consuming laboratory phenotype testing or the detection of resistance-conferring mutations in the *M. tuberculosis* genome through molecular assays or genetic sequencing.^2,3^ Although the sensitivity and specificity of these assays is high for several first and some second-line TB drugs, prediction is less accurate for other second-line antibiotics or novel agents like bedaquiline and pretomanid, which have recently become cornerstones of multi-drug resistant TB treatment. Improving the accuracy of resistance diagnosis relies on new computational approaches that can better link mutations with their functional impact on the resistance phenotype.

Two major computational strategies exist for identifying resistance-conferring variants: supervised approaches which associate genetic variation with resistance,^4^ and unsupervised approaches that search for signal of evolutionary adaptation, which can indicate resistance development. Supervised statistical methods such as genome-wide association studies and random forest classifiers have identified mutations associated with antibiotic resistance in *M. tuberculosis*,^5–9^ and machine learning approaches have built on this success to identify additional resistance-conferring variants.^10–16^ But, these approaches require a large number of phenotyped isolates (both resistant and susceptible) to make accurate predictions, which presents a major limitation, especially for recently evolved mutations. Evolution-based approaches are an alternative for finding variants associated with antibiotic resistance by analyzing their mutational frequency and phylogenetic distribution: by searching for convergent positive selection in *M. tuberculosis*, studies have found mutations in the *ald* gene associated with D-cycloserine resistance^17^, phase variation associated with virulence,^18^ and mutations involved in host-pathogen interactions that potentiate the evolution of antibiotic resistance.^19^

Both evolution-based and supervised approaches consider only the DNA or protein sequence as input. They generally assume that all sites are equally likely to affect the phenotype, and consequently many examples of a mutation are needed to infer significant effects. This assumption is not true: proteins have three-dimensional shapes and functional regions, and not every mutation is equally likely to impact function. Analyzing mutations in their three-dimensional context has uncovered hotspots of mutation in cancer,^20–24^ and provided post-hoc rationale for resistance-conferring variants in *Mycobacterium tuberculosis.*^19,25^ Protein three-dimensional structure has shown utility as an input feature for identifying resistance-conferring variants in known resistance-conferring proteins such as RpoB,^26,27^ PncA^28–30^, and AtpE.^31^While past work has sought to reannotate parts of the *M. tuberculosis* proteome with computationally predicted protein structures using older structure prediction methods,^32^ we can now infer a protein structure for nearly every protein in the proteome using AlphaFold,^33^ leading to new works examining the 3D location of mutations in known and suspected resistance-conferring proteins.^34,35^

In this paper, we use an unsupervised method to discover clustering of mutations in *M. tuberculosis* proteins, by integrating a proteome wide-structural database with mutations from over 31,000 clinical isolates. We show that mutations display statistically significant clustering in resistance-conferring genes, even in non-essential proteins such as PncA where individually rare inactivating mutations are thought to cause resistance.^36^ We identify over 450 proteins in the *M. tuberculosis* proteome that have significant clustering of mutations in their structures. Finally, we show that protein structural information provides a useful feature to predict whether variants are associated with antibiotic resistance, across all proteins with resistance variants

## Results

### Characterization of protein-modifying mutations in M. tuberculosis

We aimed to study how acquired missense variation in the *M. tuberculosis* proteome distributes in the structure of each protein. We chose to analyze the number of independent arisals of each mutation (homoplasy) rather than their population-level frequency, because analyzing the frequency of alleles in a population can be biased by oversampling of particular lineages, and by evolutionary recency,. This ensures that more recent evolutionary events are not under-represented due to lack of time to spread in the population. To accomplish this, we used a previously compiled a dataset of genomes of 31,428 isolates from the Mycobacterium tuberculosis complex (MTBC), with ancestral sequence reconstruction to determine the number of independent arisals of each mutation **(Supplementary Data 1)**.^18,19,37^ The dataset represents a diversity of MTBC isolates: with 2,815 isolates from lineage 1; 8,090 from Lineage 2; 3,398 from Lineage 3; 16,931 from Lineage 4; 98 from Lineage 5; and 96 from Lineage 6.

We filtered the 782,565 unique SNVs and 47,425 insertion/deletion mutations (INDELs) found in the original dataset to focus only missense mutations in protein coding genes, or INDELS that preserve the original translation frame. We count the number of independent arisals (homoplasy score) for each SNV and INDEL. As we are looking for positive selection that alters but does not completely ablate protein function, we do not consider frameshift mutations. After excluding mutations occurring in regions where variant calling is computationally challenging with short-read sequencing,^38,39^ we observe a total of 469,942 total unique missense mutations and 5,104 total inframe indels (**Methods, Supplementary Data 2)**. On a per-protein basis, a mean of 32.0% of sites have at least one mutation across the dataset of 31,428 isolates, with a mean of 1.4 mutations per mutated site.

We observe that proteins with the highest total number of mutations are those known to be involved in resistance to first- and second-line antibiotics **(Figure 1)**. We also observe a large number of homoplastic mutations in the protein RpoC, in which mutations can compensate for fitness loss after evolution of rifampicin resistance via mutations in RpoB.^40^ Thus, this is a marker of but not a direct cause of antibiotic resistance. Another protein with many homoplastic mutations is Cas10, a component of the CRISPR-Cas system. The vast majority of mutations events (N=1539) in this protein are due to a short inframe indel at position 3,131,469 in the H37Rv reference genome, found in over 5,000 isolates, which given the repetitive nature of the sequence region may indicate phase variation (**Supplementary Figure S1)**. Our analysis does not return ribosomal RNA genes, which are important causes of aminoglycoside resistance, because we are restricted to protein-coding genes. Finally, we note that we do not find large numbers of homoplastic mutations in genes encoding resistance to newly repurposed and introduced antibiotics linezolid, pretomanid, delamanid, bedaquiline, and clofazimine, because the majority of our dataset was sequenced before wide adoption of these drugs

**Figure 1:**
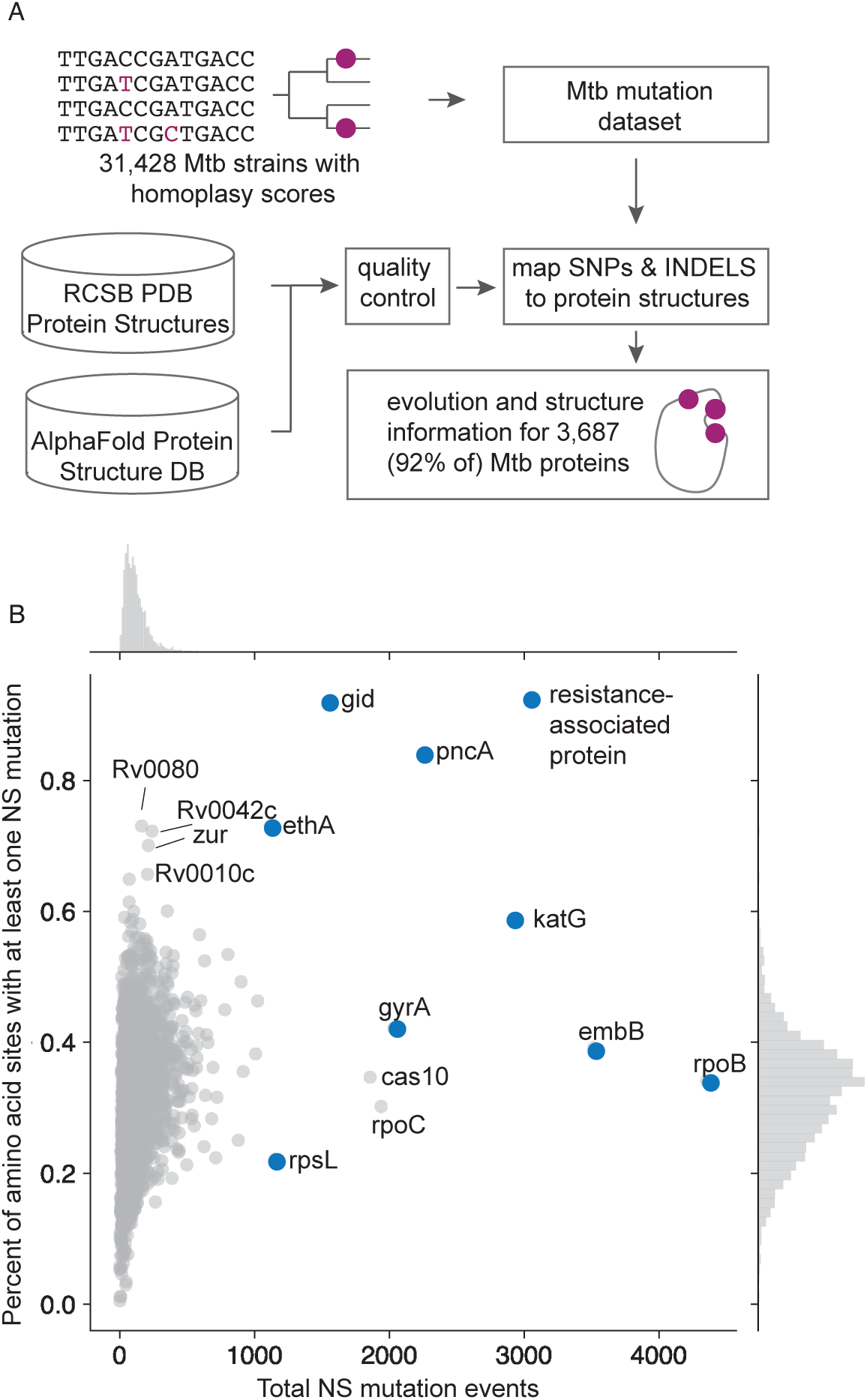
Proteins with the highest frequency of mutation events are associated with antibiotic resistance. **(A)** Workflow used to create our combined dataset of missense substitutions and inframe INDEL mutations mapped to protein 3D structures, for 92% of the *M. tuberculosis* H37Rv proteome. With a dataset of homoplastic mutations from 31,428 MTBC isolates^19,41^, we mapped these mutations to protein sequences, and 3D structures based on a combination of experimentally determined (RCSB PDB^42^) and computationally predicted (Alphafold^43^) structures. **(B)** The total number of mutation events in our dataset per protein, versus the percent of the amino acids in the protein’s structure that have been mutated at least once. Marginal histograms are displayed along both axes. Proteins with the highest frequency of mutations are those associated with resistance to antibiotics, according to the WHO catalogue of resistance-associated mutations^44^.

### Structures of the whole M. tuberculosis proteome

We identified an experimentally measured or predicted structure for each protein in the *M. tuberculosis* proteome. We used a sensitive pipeline to search the RCSB Protein structure databank (**Methods**) for structures homologous to *M. tuberculosis* proteins. We find a protein structure that represents at least 90% of the protein sequence for 34% of proteins (N = 1,371). For the remainder of proteins, we use the structure downloaded from the AlphaFold protein structure database, removing residues where the structure prediction is uncertain (pLDDT < 70). We exclude proteins with known low accuracy of variant calling (**Methods**),^32,38^ or with fewer than 30 residues represented in the filtered protein structure, for a final dataset of 3,687 proteins out of 3,996 annotated ORFs in the *M. tuberculosis* H37Rv reference proteome **(Supplementary Data 3)**.

### Antibiotic resistance proteins demonstrate mutational clustering

To calculate clustering of mutations, we use an approach from geographic analysis that calculates spatial autocorrelation between statistics, called the Getis-Ord G-statistic.^45,46^ Briefly, for each residue *i* in the protein, it calculates a *z-*score that computes if residue *i* is more proximal to highly mutated residues than would be expected by chance. This approach was previously applied to three-dimensional structure data by PIVOTAL to prioritize mutations associated with human disease.^47^

We examine the raw Getis-Ord score, which can be interpreted as a z-score, on the structures of 9 proteins in which mutations are known to confer antibiotic resistance^44^ – RpoB, EmbB, KatG, InhA, PncA, GyrA, RsmG (GidB), EthA, and Rs12 (**Methods**). For each protein, we observe a region with residue G-scores greater than 5 that are clustered in a single location **(Figure 2).** This finding is expected for essential drug target proteins where resistance-conferring mutations preserve protein function while occluding antibiotic binding, *e.g.* RpoB, EmbB, InhA, GyrA, as mutations will tend to cluster in the antibiotic binding sites of those proteins. Notably we also identify significant clustering for proteins that are not essential to the cell, where any mutation that disrupts the protein can be resistance-conferring, *e.g.* PncA. We observe that the residue G-score distribution demonstrates a periodic pattern across the gene length in which higher scores alternate with lower scores even when the high G-score residues cluster in a single protein location (**Figure 2**), a signal which emerges due to the 3D structure of the protein.

**Figure 2:**
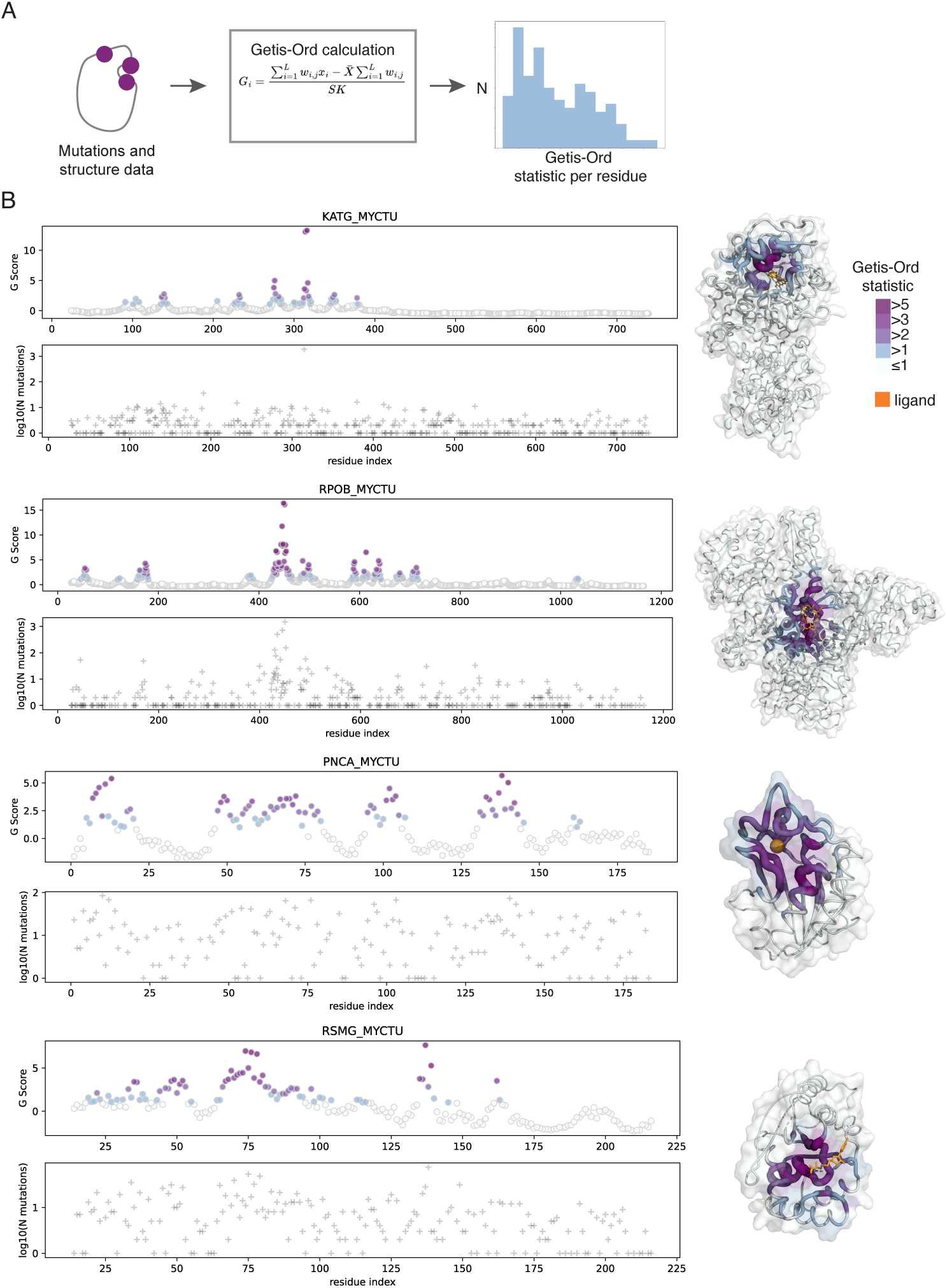
G-statistic reveals clustering of mutations in antibiotic-resistance conferring proteins. (A) Workflow to compute residue-wise Getis-Ord statistic for proteins in *M. tuberculosis.* (B) Results for proteins: KatG is shown in complex with heme (orange), PDB ID = 4C51 chain A.^48^ RpoB is shown in complex with rifampin (orange), PDB ID = 5UH6 chain C,^49^ aligned as described in **Methods.** PncA is shown in complex with Fe2+, PDB ID = 3PL1 chain A.^50^ RsmG (encoded by the *gidB* gene) structure from *Thermus thermophilus* is shown in complex with ligand adenosine monophosphate (please note the streptomycin binding site is unknown in *M. tuberculosis* and thus is not shown here), PDB ID = 3G8A chain A.^51^

### A protein-level statistic to test for mutational clustering

The G-score provides a residue-level z-score measuring three-dimensional proximity to other residues with high numbers of amino acid substitutions. We postulate that randomly accumulating substitutions would be evenly distributed in the 3-D structure of the protein. Uneven distribution of substitutions in 3D space may indicate selection; purifying selection leads to a depletion of substitutions in the hydrophobic cores of proteins, which must maintain stability,^52^ whereas positive selection may lead to mutations that cluster in protein-protein interaction interfaces or ligand binding pockets.

We evaluated five candidate protein-level statistics to quantify clustering of mutations possibly indicative of positive selection (**Methods, Supplementary Data 4**). We selected as a positive control set of proteins the nine above proteins known to be under selection for antibiotic resistance, with demonstrated residue-level clustering– RpoB, EmbB, KatG, InhA, PncA, GyrA, RsmG (GidB), EthA, and Rs12. Because each of these proteins in an outlier in terms of the overall number of mutations observed **(Figure 1B)**, we reasoned that downsampling the number of mutations would produce controls that were more similar to the average protein in the proteome. Hence for each of the nine proteins, we generate 100 positive control examples by downsampling the number of mutations observed while keeping the same relative probabilities of mutations at each site (**Figure 3, Methods**), thus preserving the signal for clustering while forcing their total number of mutations to be similar to that observed across all proteins in the proteome. We generated negative controls for the same nine proteins by simulating mutations from a uniform distribution across all residues, setting the mutation rate to the mean per-site mutation rate across the proteome, and generating 100 negative controls per protein. We find the Kolmogorov-Smirnov statistic to have the highest discrimination between positive and negative controls (95% precision, 68% recall, at significance level *p* < 0.01).

**Figure 3:**
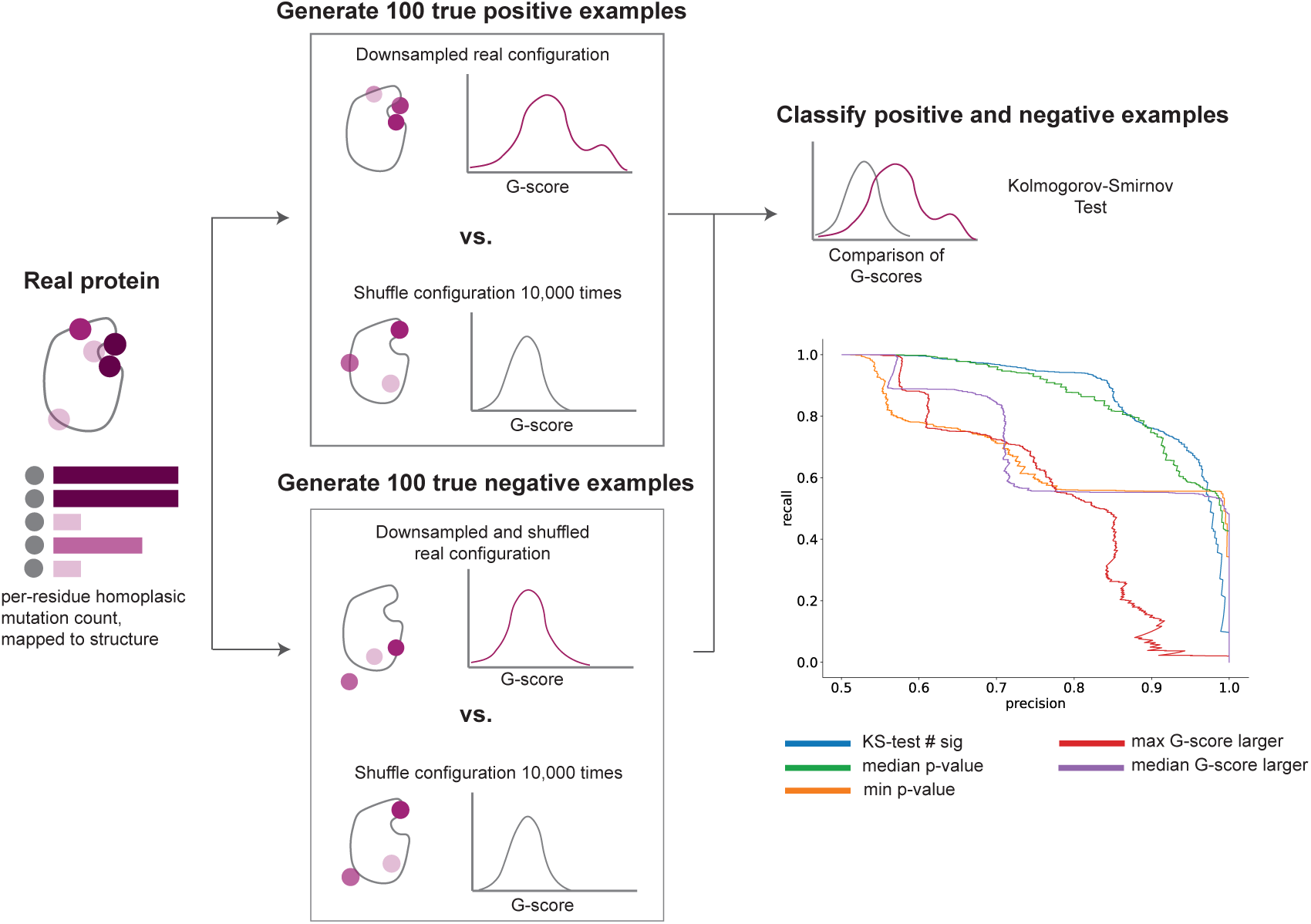
Benchmarking the ability of G-scores to find significant clustering in protein structures. The procedure for generating downsampled true positive and true negative examples from real proteins. We then test five scores for their ability to distinguish true positives and negatives.

### Significant mutational clustering across MTB proteome

We test all 3,687 proteins in the *M. tuberculosis* structure and homoplasy dataset, and identify 499 proteins with significant clustering (**Figure 4, Supplementary Data 5)**.

**Figure 4.**
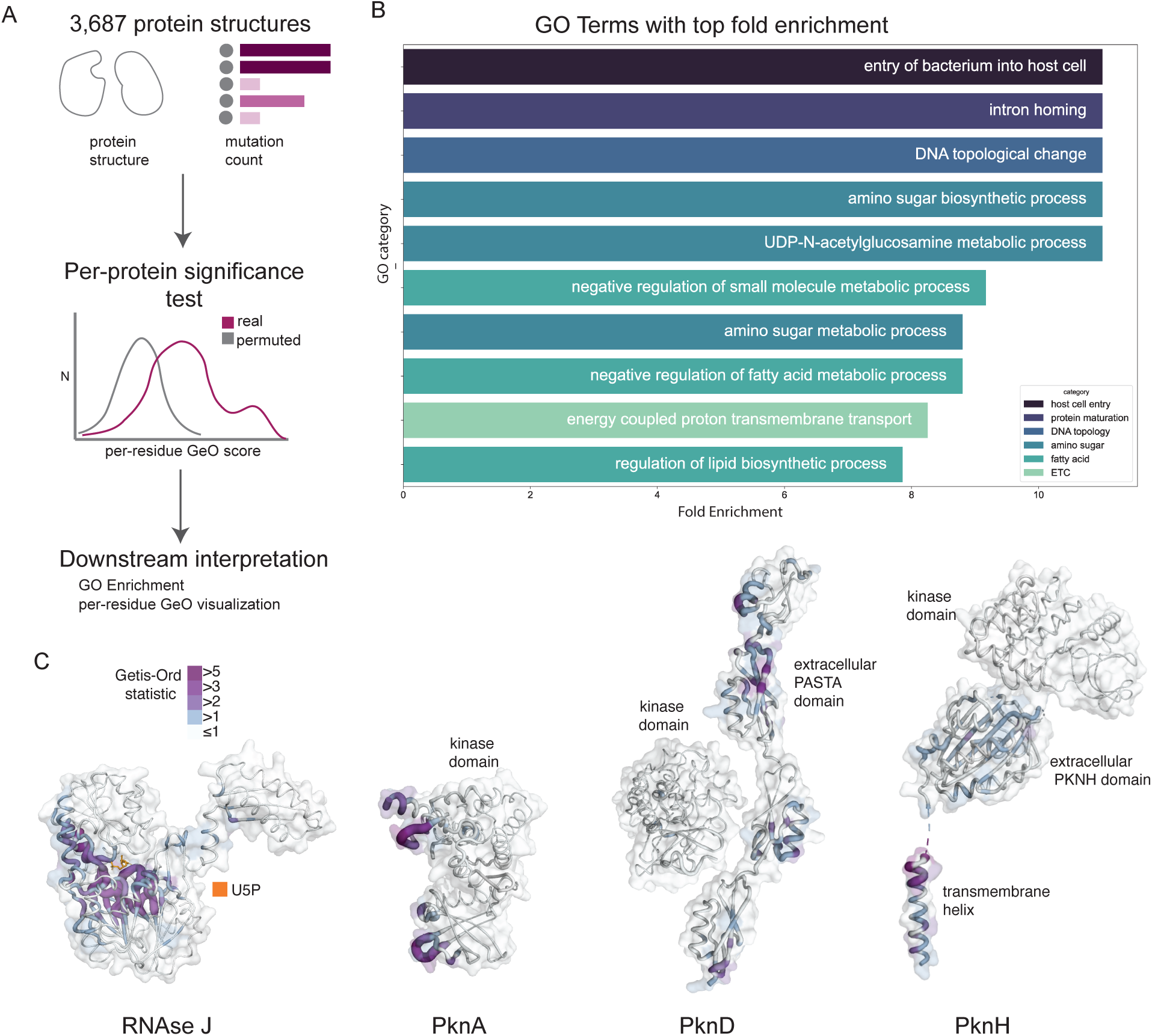
Hits of proteome-wide screen for clustering of mutations. (A) pipeline for detecting hits in 3,687 proteins. (B) GO terms with top fold enrichment (all significant at FDR < 0.05). (C) Examples of proteins with significant clustering. All structures shown are from AlphaFold with low-confidence residues filtered out (note that relative domain orientation for PknB and PknH is low confidence).

We assay whether all proteins known to cause antibiotic resistance demonstrate mutational clustering, using the WHO catalog of resistance-conferring mutations^44^. The majority of proteins with Tier-1 resistance conferring mutations (8 of 12, 67%) demonstrate significant clustering. The four proteins without clustering include TlyA and RplC, in which there is only one missense substitution known to confer resistance, and MmpR5 a transcriptional regular protein involved in resistance to the newly administered drug bedaquiline. The fourth protein is RpsL (RS12_MYCTU), whose G-score is driven by just two highly mutated residue positions, K43 and K88 (**Supplementary Figure S2**). In a small protein such as RpsL (124 amino acids), having two highly mutated residues close in 3D structure occurs with some frequency in the negative control examples.

For 90.6% (452 of 499) of the significant hits, the top G-score pair of residues are within 15 Ångstroms in 3-D space, confirming that the identified clusters are in a single location within the protein. Decreasing the distance threshold to 8 and 5 Ångstroms results in 74.7% (373) and 61.1% (305) hits whose top pairs are close in 3-D, respectively (**Supplementary Figure S3**). We find that distance between top pairs of residues does not increase with protein length, a proxy for size **(Supplementary Figure S4).** Proteins with a high number of mutations that are not spatially clustered are not significant hits. This includes Cas10, which has 1861 independent mutation events, 1489 of which occur in the same amino acid position.

We perform a Gene Ontology Enrichment analysis on the 452 hits with a close proximity top pair to understand their possible functional implications (**Methods**). After removing the less specific gene categories (with >20 genes), 54 GO categories are significantly enriched (FDR < 0.05) **(Figure 4**). The most enriched categories by fold change **(Supplementary Table 1, Supplementary Data 6)** are regulation of fatty acid metabolism and biosynthesis; UDP-N-acetylglucosamine/amino sugar metabolic processes; entry of bacterium into host; protein maturation; DNA topological change; and energy-coupled proton transport. We find clustering in the DNA topological change proteins GyrA and GyrB, both known to confer resistance to fluoroquinolones, and Top1, DNA topoisomerase I, which is not known to be associated with resistance.

A second notable GO category is composed primarily of the Pkn family of protein kinases – PknA, PknB, PknD, PknE, and PknH – thought to negatively regulate fatty acid biosynthesis. Data supports that PknA, PknB, and PknH have a role in regulating cellular permeability and thus intrinsic drug resistance, potentially through post-translational modification of proteins involved in cell wall synthesis.^53–55^ For all of the Pkn proteins found by our clustering score, the region of significant clustering is not the protein kinase domain but instead in other functional domains of the protein. For example, the significant clustering in PknB is in the extracellular domains, and in PknH is in the putative transmembrane helix and sensor domain^56^ (**Figure 4**). Therefore, we suspect that the mutations may relate to regulation of activity rather than the kinase reactions themselves.

A third notable group of proteins is involved in amino sugar metabolic processes: GlmM, GlmS, GlmU, and MurA. The related protein MurD was previously identified in a screen for targets of convergent positive selection in the *M. tuberculosis* proteome.^57^ Amino sugars are a key component of the cell wall, and thus these proteins could be under positive selection for phenotypes including intrinsic drug resistance and adaptation to the host environment.

We find significant clustering in RnJ, Ribonuclease J, which has previously been identified in GWAS for antibiotic resistance,^5^ and whose loss is known to lead to multi-drug tolerance.^58^ which has been shown to have both ribonuclease and beta lactamase activity. We find significant clustering around the AlphaFill-inferred binding site of ribonucleotides (**Methods**), which corresponds to the known active site for RNase activity (**Figure 4**).^59^

### Structure supports the prediction of resistance-conferring variants

Given that most drug resistance genes demonstrate 3-D mutational clustering, as detailed above, we tested if structural proximity to known mutations carries information on the functional impact of new mutations on phenotype. We use a previously compiled catalogue of resistance-associated mutations from *M. tuberculosis,*^9^ consisting of 583 unique missense protein-coding variants annotated as associated with resistance (R-assoc) and 58 as not associated and isolates carrying only these mutation are antibiotic susceptible (S-assoc) (**Methods**, **Supplementary Data 7**)^44^.

We ask whether protein structure information is useful at distinguishing resistance-conferring variants. In addition to the G-score computed using the homoplasy data from the ∼31,000 isolate dataset, we compute the distance in 3D between any variant and the nearest resistance-conferring variant in that protein from the catalogue. We benchmark both metrics against distance in 1D (on the protein sequence). We trained a logistic regression classifier to predict whether a mutation is resistance associated (R-assoc, *vs.* S-assoc, susceptibility associated), using a 70-30 train-test split of the cataloged mutations. 3D structural proximity alone best predicted resistance association (F1 score of 94.6%, **Table 1, Methods)**. Prediction from 1D proximity alone is less accurate at F1 of 92.8%, and prediction from G-score had the lowest F1 at 80.8%. The relatively strong performance of 1D proximity prediction is due to the fact that most, but not all, mutations in our dataset that have close 3D proximity to a mutated residue also have close 1D proximity.

**Table 1:** Performance of classification models on predicting whether mutations are R-conferring from the mutation catalog. Proximity-1D was trained using just the distance in primary sequence to the nearest known R mutation, Proximity-3D was trained using the distance in 3D to the nearest known R mutation, and G-score was trained using the G-score calculated in this manuscript.

| Feature Set | F1 | Precision/PPV | Sensitivity |
| --- | --- | --- | --- |
| 3D-Proximity | 94.6 | 97.5 | 91.9 |
| 1D-Proximity | 92.8 | 96.3 | 89.5 |
| G-score | 80.8 | 98.3 | 68.6 |

While the G-score prediction has the lowest F1 score and sensitivity, it has the highest precision (98.3%). This may be because the G-score is a combination of structural information and mutational information, as a residue will only have a high G-score if it is proximal to at least one residue with a high number of mutations (presumably due to being causal of resistance). This means that the G-score model is more conservative at calling R-assoc variants as truly R-assoc.

## Discussion

In this paper, we demonstrate that mutations cluster in three dimensions in the structure of proteins known to confer antibiotic resistance, through a novel application of the Getis-Ord statistic to evolutionary sequence data from clinical tuberculosis isolates. We apply our method to all proteins in the *M. tuberculosis* H37Rv proteome, using homoplasy data generated from a dataset of over 30,000 *M. tuberculosis* complex isolates. We find significant clustering of mutations in over 450 proteins, including eight known resistance-conferring proteins, and in pathways thought to be important for pathogenesis and antibiotic resistance.

We observe spatial clustering of mutations in known resistance-conferring proteins. For proteins like RpoB, where mutations are known to cluster in the rifampicin binding site, this serves as an internal control for our method. However, for proteins where mutations that lead to loss of function are known to cause resistance, such as PncA and RsmG (GidB), it is not necessarily expected to find clustering of mutations. We suspect that the observed clustering is due to mutations in a certain region of the protein being more likely to cause loss of function.

When running our analysis on the entire proteome, we find a significant enrichment for hits in proteins involved in regulation of fatty acid biosynthetic processes, and in amino sugar metabolism. Both of these pathways are related to cell envelope synthesis, which may be a response to antibiotic pressure or an adaptation to host environments. We also find hits to the proteins MycP1, P2, and P3, which are secreted from the cell and thought to be involved in host cell entry. We chose to apply our approach to homoplasy data, not allele frequency data, to ensure that we were not biased to overweight more ancient mutations in our analysis. Thus, we believe significant Getis-Ord score clustering indicates protein hotspots of positive and/or diversifying selection, where *M. tuberculosis* is adapting to antibiotic pressure and host environments.

Our variant effect prediction results support that structural information is a meaningful feature for predicting whether specific genetic variants confer antibiotic resistance. Distance to ligand binding sites have previously been shown to be a meaningful feature for classification of resistance in the proteins RpoB,^26,27^ PncA,^28^ and AtpE,^31^ and our work extends this to all proteins with known resistance-conferring mutations. These results on resistance variant classification how potential methods for expanding the catalog of known resistance-conferring mutations. In addition, recent efforts to predict *M. tuberculosis* antibiotic resistance from sequence data have suggested that our ability to predict antibiotic resistance from sequence alone may be reaching a plateau, and that more data and different data modalities, not more innovative model architectures, are needed in order to further improve predictions.^12^ While we do not suggest that our current model be used in clinical settings, we envision structure being added as an additional feature in future work to predict organismal resistance phenotypes from sequences, as it has already been successfully used in resistance variant classification.^34,35^

One drawback of our approach is that our random shuffling procedure implicitly assumes that all sites in a protein are equally likely to mutate. This may pose a problem for proteins with conserved hydrophobic cores, or multidomain proteins where one domain is more conserved than the other, which could manifest in apparent higher or lower rates of mutation in one region of the protein. We account for this bias when we compute the distance between the top-scoring pair of residues in each protein, finding that for over 90% of the proteomic hits, those two residues are within 15 Ångstroms of one another, indicating that the patch of high scoring residues are in a single location on the protein, not spread throughout a surface or domain.

However, we cannot entirely rule out the effects of differing mutation rates throughout a protein. Future work could consider simulating the accumulation of substitutions according to their predicted effects on protein stability, to better reflect a real-world evolutionary scenario.

The research community has long sought to determine the genetic basis of antibiotic resistance in *M. tuberculosis,* using both supervised methods that rely on labelled data, and unsupervised methods that seek to find patterns of positive selection indicative of antibiotic resistance. The availability of predicted protein structures has opened a new avenue for understanding the effects of mutations in microbial genomes, and in *M. tuberculosis* specifically. We hypothesize that protein structural proximity is a useful feature because it captures how likely any given pair of mutations are to have a similar phenotypic effect – mutations in the same functional regions of proteins are more likely to cause the same effect. We show that protein 3D structure can be used to classify resistance-conferring variants across all proteins in a supervised framework, and used in an unsupervised fashion to discover targets of positive selection in the *M. tuberculosis* genome.

## Methods

### Mycobacterium tuberculosis mutation dataset

We use a dataset of SNPs and indels found in *M. tuberculosis* isolates, mapped to their occurrences on a phylogenetic tree, published by previous work.^18,37^ This dataset was constructed using a previously validated pipeline for calling variants in *M. tuberculosis,* by mapping reads to the H37Rv reference genome using BWA-MEM v0.7.17^60,61^ after trimming and filtering with PRINSEQ v0.20.4,^62^ contaminant removal with Kraken v0.10.6,^63^ and duplicate read removal with Picard v2.9.2. All isolates were required to have at least 95% of bases in the reference genome with at least 10x coverage. Variant calling was performed with Pilon,^64^ and additional quality control filters were applied to ensure accurate allele calls. The resulting dataset was two matrices: a list of all positions found to have a SNP in any isolate, relative to the H37Rv reference (N=782,565), and a list of all positions found to have an insertion or deletion in any isolate, relative to the H37Rv reference (N=47,425).^18^

Phylogeny and ancestral sequence reconstruction were performed by Vargas et al as described in a previous publication.^18^ In their procedure, phylogenies were constructed separately for sub-lineages (L1, L2, L3, L4A, L4B, L4C, L5, L6) for computational feasibility. Phylogenies were constructed from concatenated SNP data using IQTree.^65^ SNPPar^66^ was used to reconstruct SNVs, and the method previously published by Vargas *et al.* was used to reconstruct INDELs.^41^

### Searching PDB structure database

We search the RCSB Protein Data Bank structure database (download date: Feb 5, 2021) for experimentally determined structures with sequence similarity to the Uniprot Proteome of Mycobacterium tuberculosis (ID = UP000001584). In order to execute a sensitive search for homologous structures, we built a modified version of the EVcouplings pipeline,^67^ which operates in two stages: first, we construct sequence alignments against the Uniprot sequence database (download date Feb 5, 2021) at bitscores of 0.1, 0.2, and 0.3 times the query sequence length, using jackhmmer from hmm-suite v3.1 with five iterations^68^. Second, we use the hmmbuild and hmmsearch tools of hmm-suite to search the RCSB PDB sequence database using the constructed alignments.

### Selecting structure hits

We then aim to select high-coverage protein structures from the experimental structure database to represent the *M. tuberculosis* proteins. We define coverage as whether an amino acid residue in the *M. tuberculosis* protein is represented by a resolved amino acid in the protein structure. Of the 3993 proteins in the proteome, 2984 (75%) have any hits to the structure database. 1509 (38%) of these hits have at least 90% coverage. In cases where more than one hit had 90% coverage, we select the representative hit in the following way: if a protein structure is from *M. tuberculosis*, we select that structure, otherwise we select the hit with the lowest e-value.

### Using AlphaFold database

Predicted structures for the Mycobacterium tuberculosis reference proteome (ID = UP000001584) were downloaded from the AlphaFold Protein Structure Database on Feb 15, 2023. Residues with pLDDT < 70 were removed from each structure.

### Preparing protein structure data for analysis

We exclude from consideration proteins with documented difficulties in mutation calling using short-read sequencing. Using data from Marin *et al*^38^, proteins with <90% mean empirical base pair recall (across all residues in the protein) or <90% mean residue mappability were removed from consideration. Proteins with fewer than 30 residues with resolved structure coordinates were removed, for a final total of 3687 proteins, 1350 of which are experimentally determined structures and the remaining 2337 of which are from AlphaFold.

### Preparing mutations for analysis

For SNP mutations, we consider all missense SNPs that occur at least once in our *M. tuberculosis* dataset, for a total of 476,607 unique polymorphisms. For insertion and deletion (indel) mutations, we exclude frameshift mutations as these are likely to ablate protein function and thus are not analogous to missense mutations. We observe 5,678 unique in-frame indels. Combining the SNP and inframe indel mutations, we observe 483,117 unique mutations. To account for possible artifacts from short-read sequencing base calling errors, we exclude mutations in positions designated as blindspots by Modlin *et al* or with empirical base-pair recall <90%,^38,39^ as well as all mutations occurring in proteins with <90% mean empirical base-pair recall or <90% mean mappability.^22^ After filtering, we have 475,046 unique polymorphisms for downstream analysis (469,942 total unique missense mutations and 5,104 INDELs) (**Supplementary Data 2)**.

### Merging mutation and structure data

Upon merging our structure dataset with our mutation dataset, 335,224 mutations can be mapped to a residue in a protein with high-quality structure information. We exclude proteins with fewer than two mutations mapped to their structure from the clustering calculation. Indel mutations were treated as occurring at the first position in the sequence. We proceed with clustering analysis on 3657 proteins with mutations mapped.

### Computing inter-residue distances

The EVcouplings Python package was used to compute the distance between wild-type amino acid residues in all protein structures.^67^ The package calculates the distance between all heavy (non-hydrogen) atoms in residue *i* and residue *j*, then returns the minimum of those distances.

### Computing the Getis-Ord score for clustering of homoplastic mutations

Our implementation of the Getis-Ord score for protein three-dimensional structure was inspired by PIVOTAL, which applied the Getis-Ord score to clustering of mutations in human disease.^45–47^ The two values input to the Getis-Ord statistic computation are a per-residue score **x,** here the per-amino acid homoplasy score, and a weight matrix *W* that contains the inverse of the inter-residue distances computed from the wild-type amino acids. **x** is an L x 1 vector where the entry *x*_!_ contains the homoplasy score for the i-th residue in the protein, and **W** is an L x L matrix with entries defined as:

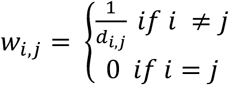

Where *d_i,j_* is the minimum inter-residue atomic distance, in Å, between i and j. Thus, residues that are closer in three dimensions will have higher weights.

The Getis-Ord statistic for residue i is calculated as:

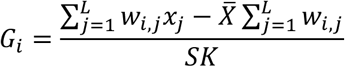

Where *X̅* is the mean of **x,** and:

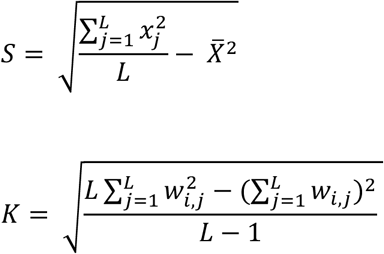

### Preparing GeO score calibration data

We sought to generate a dataset of positive controls – proteins known to have clustering of mutations – with a frequency of mutation similar to the average protein in our dataset (mean of 1.4 mutations per site). For this, we used the nine proteins known to be involved in antibiotic resistance with demonstrated clustering of mutations (RpoB, EmbB, KatG, InhA, PncA, GyrA, RsmG (GidB), EthA, and Rs12). For each of these nine proteins, we generate 100 positive control examples.

Because the total number of mutations in a site, as well as the 3D configuration of sites, contributes to the G-score, we chose to downsample these proteins to generate positive controls that are more similar to other proteins in the total number of mutations. To generate positive control examples, we build an empirical distribution based on the observed number of mutations per site in the protein, divided by the total mutations. We add a pseudocount of one to all residues with zero mutations. We then sample M mutations from the empirical mutation distribution, where M = 0.42 times the number of residues in the protein with structure data, because each site is mutated on average 0.42 times. This sampling strategy preserves the relative frequencies of each mutation, and their 3D location, while greatly reducing the number of total mutations observed in a protein. We generate the negative controls by sampling from a uniform distribution over all residues in a protein with structure data. Note that we do not re-compute inter-residue distances when simulating mutations in an amino acid, as the distances used as input to GeO score are the wild-type inter-residue distances.

Then, we build a heuristic based on the Getis-Ord score to determine if we observe significant clustering in a protein. For each control example, we perform 10,000 random permutations of the mutations, shuffling their configuration in space but keeping the number of mutation events the same. For positive controls, we expect the real configuration of mutations to produce a significantly different configuration of G scores than random permutations. For negative controls, we do not expect the real configuration of mutations to produce a significantly different configuration of G scores than random permutations.

Next, for each generated control, we test the ability of various scores to distinguish between the real example and a random reshuffling. By comparing the true distribution with the empirical random distribution, we compute a p-value for whether the overall distribution of Getis-Ord statistics for residues in the protein is different than the one expected by chance. We compare six scores on the basis of their precision recall curves for recovering the positive controls. The first three scores are based on comparing the p-value of the Kolmogorov-Smirnov test when comparing the real distribution to 10,000 random shuffles: the number of significant p-values (*p* < 0.01), the smallest p-value, and the mean p-value. The other two features are based on the raw G-score for the real distribution vs. the 10,000 random shuffles: how often the max G-score is greater for the real distribution, and how often the median G-score is greater for the real distribution.

### Running on whole proteome

We then run the G-score calculation on the whole proteome using the above pipeline. We apply our 95% precision threshold to find 451 initial hits.

### GO Enrichment Analysis

We test whether our gene hits are enriched in particular GO functional categories.^69,70^ We use the online GO enrichment tool (https://geneontology.org/) to search for enriched biological processes among our 452 hit proteins. We downloaded the .json file from the GO enrichment tool, and filtered for GO categories with fewer than 20 members in the *M. tuberculosis* H37Rv refence genome, to remove overly broad categories like “biological process,” and terms with identical constituent genes. We retained categories with FDR < 0.05, for a total of 37 enriched terms.

### AlphaFill

We downloaded hits from the AlphaFill v1 database (access date: 7 October 2024) to find potential ligand-binding locations in our proteins. For RnJ, two ligands are in proximity of the high GeO score regions: “U5P” and “C5P”, uridine-5’-monophosphate and cytidine-5’-monophosphate.

### Parsing the WHO mutation catalog

The World Health Organization (WHO) Mutation Catalogue maintains a list of over 30,000 unique variants observed in *M. tuberculosis* genomes and whether those mutations are diagnostic of antibiotic resistance against 13 antibiotics^71^. Mutations are graded with five confidence categories: (1) Associated with Resistance, (2) Associated with Resistance - Interim, (3) Uncertain Significance, (4) Not Associated with Resistance - Interim, and (5) Not Associated with Resistance. Kulkarni *et al.* provided an update to this catalog which increased the number of labelled variants using regression-based grading based on frequency of mutations in resistant and susceptible isolates.^9^

We limit our analysis to the variants observed in the following 15 protein-coding genes: *Rv0678, atpE, ddn, embB, ethA, gid, gyrA, gyrB, inhA, katG, pncA, rplC, rpoB, rpsL, tlyA.* After removing duplicates – because multiple genomic variants can cause the same missense mutation – we were left with 9365 variants, composed mostly of uncertain variants. After filtering for variants with structure information, we have 583 R-assoc (category 1 or 2), and 58 as S-assoc (category 4 or 5).

### Fitting a classifier on the WHO mutation catalog

For each of 9365 unique missense variants in the catalogue,^9^ we extracted the following features: The minimum coordinate difference to the nearest non-self R variant along the amino-acid sequence (“1D proximity”). The inter-atomic distance to the nearest non-self R variant in Ångstroms (“3D proximity”), the G-score of the residue as computed in our previous analysis. 3D proximity was calculated using the EVcouplings Python package.^67^

We used a 70-30 train-test split to construct a dataset. We employed weighted sampling to address the class imbalance between resistance-conferring and non-resistance-conferring variants. We employed a logistic regression classifier in scikit-learn, and hyperparameters were tuned by grid search.^72^ Models were evaluated for F1-Score, precision, and recall. To select an optimal model threshold, we choose the threshold which maximizes the sum of sensitivity and specificity.^73^

## Acknowledgements

We thank members of the Farhat lab at Harvard Medical School and the SAGE lab at University of Massachusetts Amherst for valuable discussion about the project. Computational resources and support were provided by the Orchestra High Performance Compute Cluster at Harvard Medical School, which is funded by the NIH (NCRR 1S10RR028832-01). A.G.G. was supported by NIH/NIAID F32AI161793. R.V.J. was supported by the National Science Foundation Graduate Research Fellowship under Grant No. DGE1745303.

## Author Contributions

A.G.G. and M.R.F. conceived the study and designed the analyses. A.G.G. implemented the mutation clustering code. M.T. implemented the resistance mutation classification code. A.G.G., M.T., and M.R.F. interpreted the data and results. R.V.J contributed data and provided important discussions of results. M.R.F. and A.G.G. supervised the research. A.G.G., M.T., and M.R.F. wrote the manuscript with input from all authors.

## Data Availability

Code is available on GitHub at https://github.com/aggreen/MTB_Mut_Clust, and the modified EVcouplings pipeline for structure search is found at https://github.com/aggreen/EVcouplings/tree/feature/structure_finder. All strains used in our analyses are publicly available, and the raw read data are available for download from the NCBI using accession codes found in the isolate annotation table. WHO mutation classification data 2^nd^ edition is publicly available here: https://www.who.int/publications/i/item/9789240082410

## References

1. World health Organization. Global Tuberculosis Report 2021. (2021).

2. Global Tuberculosis Programme, W. H. (hq). WHO Consolidated Guidelines on Tuberculosis. Module 3: Diagnosis - Rapid Diagnostics for Tuberculosis Detection 2021 Update. (2021).

3. Lange, C. et al. Drug-resistant tuberculosis: An update on disease burden, diagnosis and treatment. Respirology 23, 656–673 (2018).

4. Bush, W. S. & Moore, J. H. Chapter 11: Genome-wide association studies. PLoS Comput Biol 8, e1002822 (2012).

5. Farhat, M. R. et al. GWAS for quantitative resistance phenotypes in Mycobacterium tuberculosis reveals resistance genes and regulatory regions. Nat Commun 10, 2128 (2019).

6. Gröschel, M. I. et al. GenTB: A user-friendly genome-based predictor for tuberculosis resistance powered by machine learning. Genome Med 13, 138 (2021).

7. Conkle-Gutierrez, D. et al. Distribution of Common and Rare Genetic Markers of Second-Line-Injectable-Drug Resistance in Mycobacterium tuberculosis Revealed by a Genome-Wide Association Study. Antimicrob. Agents Chemother. 66, e02075–21 (2022).

8. Coll, F. et al. Genome-wide analysis of multi- and extensively drug-resistant Mycobacterium tuberculosis. Nat Genet 50, 307–316 (2018).

9. Kulkarni, S. G. et al. Regression for accurate and sensitive grading of mutations diagnostic of antibiotic resistance in Mycobacterium tuberculosis. 2024.07.01.24309598 Preprint at 10.1101/2024.07.01.24309598 (2024).

10. Green, A. G. et al. A convolutional neural network highlights mutations relevant to antimicrobial resistance in Mycobacterium tuberculosis. Nat Commun 13, 3817 (2022).

11. Chen, M. L. et al. Beyond multidrug resistance: Leveraging rare variants with machine and statistical learning models in Mycobacterium tuberculosis resistance prediction. EBioMedicine 43, 356–369 (2019).

12. Wang, Y. et al. TB-DROP: deep learning-based drug resistance prediction of Mycobacterium tuberculosis utilizing whole genome mutations. BMC Genomics 25, 167 (2024).

13. Pruthi, S. S. et al. Leveraging large-scale Mycobacterium tuberculosis whole genome sequence data to characterise drug-resistant mutations using machine learning and statistical approaches. Sci. Rep. 14, 27091 (2024).

14. Serajian, M. et al. Scalable de novo classification of antibiotic resistance of Mycobacterium tuberculosis. Bioinformatics 40, i39–i47 (2024).

15. Kulkarni, S. G. et al. Convolutional neural networks quantify antibiotic resistance in Mycobacterium tuberculosis with diagnostic grade accuracy and predict treatment response. 2025.08.05.25333066 Preprint at 10.1101/2025.08.05.25333066 (2025).

16. Pal, A. & Mohanty, D. Machine learning-based approach for identification of new resistance associated mutations from whole genome sequences of Mycobacterium tuberculosis. Bioinforma. Adv. 5, vbaf050 (2025).

17. Desjardins, C. A. et al. Genomic and functional analyses of Mycobacterium tuberculosis strains implicate ald in D-cycloserine resistance. Nat Genet 48, 544–551 (2016).

18. Vargas, R., et al. Phase variation as a major mechanism of adaptation in Mycobacterium tuberculosis complex. Proc. Natl. Acad. Sci. 120, e2301394120 (2023).

19. Green, A. G. et al. Analysis of Genome-Wide Mutational Dependence in Naturally Evolving Mycobacterium tuberculosis Populations. Mol. Biol. Evol. 40, msad131 (2023).

20. Miller, M. L. et al. Pan-Cancer Analysis of Mutation Hotspots in Protein Domains. Cell Syst. 1, 197–209 (2015).

21. Gao, J. et al. 3D clusters of somatic mutations in cancer reveal numerous rare mutations as functional targets. Genome Med 9, 4 (2017).

22. Kamburov, A., et al. Comprehensive assessment of cancer missense mutation clustering in protein structures. Proc. Natl. Acad. Sci. 112, E5486–E5495 (2015).

23. Meyer, M. J. et al. mutation3D: Cancer Gene Prediction Through Atomic Clustering of Coding Variants in the Structural Proteome. Hum. Mutat. 37, 447–456 (2016).

24. Niu, B. et al. Protein-structure-guided discovery of functional mutations across 19 cancer types. Nat. Genet. 48, 827–837 (2016).

25. Phelan, J. et al. Mycobacterium tuberculosis whole genome sequencing and protein structure modelling provides insights into anti-tuberculosis drug resistance. BMC Med 14, (2016).

26. Lynch, C. I., Adlard, D. & Fowler, P. W. Predicting rifampicin resistance in Mycobacterium tuberculosis using machine learning informed by protein structural and chemical features. ERJ Open Res. 11, 00952–02024 (2025).

27. Portelli, S. et al. Prediction of rifampicin resistance beyond the RRDR using structure-based machine learning approaches. Sci. Rep. 10, 18120 (2020).

28. Karmakar, M., Rodrigues, C. H. M., Horan, K., Denholm, J. T. & Ascher, D. B. Structure guided prediction of Pyrazinamide resistance mutations in pncA. Sci. Rep. 10, 1875 (2020).

29. Carter, J. J., et al. Prediction of pyrazinamide resistance in Mycobacterium tuberculosis using structure-based machine-learning approaches. JAC-Antimicrob. Resist. 6, dlae037 (2024).

30. Dissanayake, D., Brunner, V., Adlard, D., Morrone, J. A. & Fowler, P. W. Predicting pyrazinamide resistance in Mycobacterium tuberculosis using a graph convolutional network. 2025.10.28.685176 Preprint at 10.1101/2025.10.28.685176 (2026).

31. Karmakar, M. et al. Empirical ways to identify novel Bedaquiline resistance mutations in AtpE. PLOS ONE 14, e0217169 (2019).

32. Modlin, S. J. et al. Structure-Aware Mycobacterium tuberculosis Functional Annotation Uncloaks Resistance, Metabolic, and Virulence Genes. mSystems 6, e00673–21 (2021).

33. Jumper, J. et al. Highly accurate protein structure prediction with AlphaFold. Nature 596, 583–589 (2021).

34. Pal, A., Pal, S., Mahapatra, S., Pandey, A. & Mohanty, D. *In silico* analysis of the functional implications of drug resistance associated mutations in *Mycobacterium tuberculosis*. Comput. Struct. Biotechnol. J. 27, 5425–5440 (2025).

35. Wood, J. J., Portelli, S., Ascher, D. B. & Furnham, N. Risk-Based Prediction of Novel AMR Variants Using Protein Language Models. 2025.09.12.672331 Preprint at 10.1101/2025.09.12.672331 (2026).

36. Farhat, M. R. et al. Genetic Determinants of Drug Resistance in Mycobacterium tuberculosis and Their Diagnostic Value. Am J Respir Crit Care Med 194, 621–630 (2016).

37. Vargas, R. et al. Role of Epistasis in Amikacin, Kanamycin, Bedaquiline, and Clofazimine Resistance in Mycobacterium tuberculosis Complex. Antimicrob. Agents Chemother. 65, 10.1128/aac.01164-21 (2021).

38. Marin, M. et al. Benchmarking the empirical accuracy of short-read sequencing across the M. tuberculosis genome. Bioinformatics (2022).

39. Modlin, S. J. et al. Exact mapping of Illumina blind spots in the Mycobacterium tuberculosis genome reveals platform-wide and workflow-specific biases. Microb Genom 7, (2021).

40. Comas, I. et al. Whole-genome sequencing of rifampicin-resistant Mycobacterium tuberculosis strains identifies compensatory mutations in RNA polymerase genes. Nat Genet 44, 106–110 (2011).

41. Vargas, R., et al. Phase variation as a major mechanism of adaptation in Mycobacterium tuberculosis complex. bioRxiv (2022).

42. Berman, H. M. et al. The protein data bank. Nucleic Acids Res 28, 235–242 (2000).

43. Varadi, M. et al. AlphaFold Protein Structure Database: massively expanding the structural coverage of protein-sequence space with high-accuracy models. Nucleic Acids Res 50, D439–D444 (2022).

44. Walker, T. M. et al. The 2021 WHO catalogue of Mycobacterium tuberculosis complex mutations associated with drug resistance: A genotypic analysis. Lancet Microbe 3, e265–e273 (2022).

45. Getis, A. & Ord, J. K. The Analysis of Spatial Association by Use of Distance Statistics. Geogr. Anal. 24, 189–206 (1992).

46. Ord, J. K. & Getis, A. Local Spatial Autocorrelation Statistics: Distributional Issues and an Application. Geogr. Anal. 27, 286–306 (1995).

47. Liang, S., Mort, M., Stenson, P. D., Cooper, D. N. & Yu, H. PIVOTAL: Prioritizing variants of uncertain significance with spatial genomic patterns in the 3D proteome. 2020.06.04.135103 Preprint at 10.1101/2020.06.04.135103 (2021).

48. Zhao, X., Hersleth, H.-P., Zhu, J., Andersson, K. K. & Magliozzo, R. S. Access channel residues Ser315 and Asp137 in Mycobacterium tuberculosis catalase-peroxidase (KatG) control peroxidatic activation of the pro-drug isoniazid. Chem. Commun. 49, 11650–11652 (2013).

49. Lin, W. et al. Structural Basis of *Mycobacterium tuberculosis* Transcription and Transcription Inhibition. Mol. Cell 66, 169–179.e8 (2017).

50. Petrella, S. et al. Crystal structure of the pyrazinamidase of Mycobacterium tuberculosis: insights into natural and acquired resistance to pyrazinamide. PLoS One 6, e15785 (2011).

51. Gregory, S. T. et al. Structural and functional studies of the Thermus thermophilus 16S rRNA methyltransferase RsmG. RNA 15, 1693–1704 (2009).

52. Echave, J., Spielman, S. J. & Wilke, C. O. Causes of evolutionary rate variation among protein sites. Nat Rev Genet 17, 109–121 (2016).

53. Sun, M., Ge, S. & Li, Z. The Role of Phosphorylation and Acylation in the Regulation of Drug Resistance in Mycobacterium tuberculosis. Biomedicines 10, 2592 (2022).

54. Zeng, J. et al. Protein kinases PknA and PknB independently and coordinately regulate essential Mycobacterium tuberculosis physiologies and antimicrobial susceptibility. PLoS Pathog. 16, e1008452 (2020).

55. Sharma, K. et al. Transcriptional Control of the Mycobacterial embCAB Operon by PknH through a Regulatory Protein, EmbR, In Vivo. J. Bacteriol. 188, 2936–2944 (2006).

56. Cavazos, A., Prigozhin, D. M. & Alber, T. Structure of the Sensor Domain of *Mycobacterium tuberculosis* PknH Receptor Kinase Reveals a Conserved Binding Cleft. J. Mol. Biol. 422, 488–494 (2012).

57. Farhat, M. R. et al. Genomic analysis identifies targets of convergent positive selection in drug-resistant Mycobacterium tuberculosis. Nat Genet 45, 1183–1189 (2013).

58. Martini, M. C. et al. Loss of RNase J leads to multi-drug tolerance and accumulation of highly structured mRNA fragments in Mycobacterium tuberculosis. PLoS Pathog. 18, e1010705 (2022).

59. Bao, L. et al. Structural insights into RNase J that plays an essential role in Mycobacterium tuberculosis RNA metabolism. Nat. Commun. 14, 2280 (2023).

60. Li, H. Aligning sequence reads, clone sequences and assembly contigs with BWA-MEM. (2013).

61. Li, H. & Durbin, R. Fast and accurate short read alignment with Burrows–Wheeler transform. Bioinformatics 25, 1754–1760 (2009).

62. Schmieder, R. & Edwards, R. Quality control and preprocessing of metagenomic datasets. Bioinformatics 27, 863–864 (2011).

63. Wood, D. E. & Salzberg, S. L. Kraken: ultrafast metagenomic sequence classification using exact alignments. Genome Biol 15, R46 (2014).

64. Walker, B. J. et al. Pilon: an integrated tool for comprehensive microbial variant detection and genome assembly improvement. PLoS One 9, e112963 (2014).

65. Nguyen, L.-T., Schmidt, H. A., von Haeseler, A. & Minh, B. Q. IQ-TREE: A Fast and Effective Stochastic Algorithm for Estimating Maximum-Likelihood Phylogenies. 10.1093/molbev/msu300.

66. Edwards, D. J., Duchene, S., Pope, B. & Holt, K. E. SNPPar: identifying convergent evolution and other homoplasies from microbial whole-genome alignments. *Microb*. Genomics 7, 000694 (2021).

67. Hopf, T. A. et al. The EVcouplings Python framework for coevolutionary sequence analysis. Bioinformatics 35, 1582–1584 (2019).

68. HMMER. http://hmmer.org/.

69. Ashburner, M. et al. Gene ontology: tool for the unification of biology. The Gene Ontology Consortium. Nat Genet 25, 25–29 (2000).

70. The Gene Ontology Consortium et al. The Gene Ontology knowledgebase in 2023. Genetics 224, iyad031 (2023).

71. Catalogue of mutations in Mycobacterium tuberculosis complex and their association with drug resistance, 2nd ed. https://www.who.int/publications/i/item/9789240082410.

72. Pedregosa, F. et al. Scikit-learn: Machine Learning in Python. J Mach Learn Res 12, 2825–2830 (2011).

73. Ruopp, M. D., Perkins, N. J., Whitcomb, B. W. & Schisterman, E. F. Youden Index and Optimal Cut-Point Estimated from Observations Affected by a Lower Limit of Detection. Biom. J. Biom. Z. 50, 419–430 (2008).

