## Supplementary Figures and Tables for "The structural context of mutations in proteins predicts their effect on antibiotic resistance"

**Supporting Information**

**Figure S1: Homoplastic inframe insertion at position 3131469 in the Cas10 gene (Rv2823c).** Screenshot from Mycobrowser (https://mycobrowser.epfl.ch/) of relevant genomic region beginning at 3131469. Table showing the number of mutation events per lineage and total in the dataset, as well as total number of isolates with the alternate allele. Note that the inserted sequence is similar to but not exactly the same as the H37Rv reference sequence at that location, and that similar motifs recur throughout the sequence region.

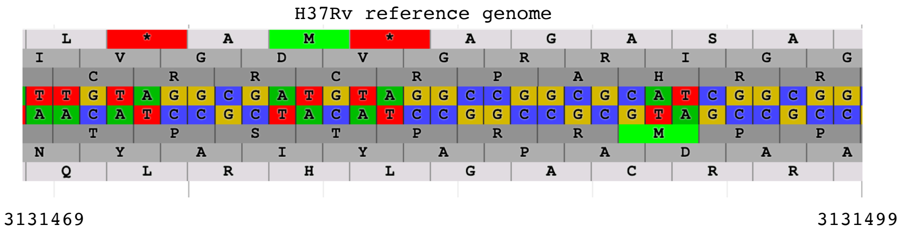


| | | | | | N mutation events per lineage | | | | | | | | | |
|---|---|---|---|---|---|---|---|---|---|---|---|---|---|---|
| Position | Ref | Alternate | Gene Name | INDEL type | L1 | L2 | L3 | L4A | L4B | L4C | L5 | L6 | total | N isolates with alternate allele |
| 3131469 | T | TTGTCGGCGA | Rv2823c | inframe | 158 | 377 | 183 | 351 | 367 | 88 | 5 | 5 | 1534 | 5093 |

**Figure S2: Mutations and GeO clustering for RpsL (RS12_MYCTU).** Two residues, shown in orange (K43 and K88) are highly mutated in the protein RpsL, leading to significant G-score clustering in that region of the protein.

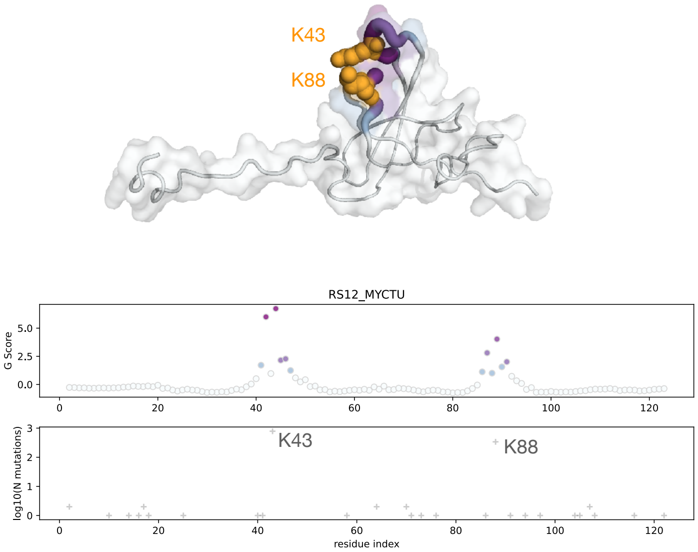

**Supplementary Figure S3: Distance between top pairs of high G-score residues in proteins with significant clustering.** Of the 499 proteins with significant hits, we analyze what number are still significant after filtering to ensure that the minimum inter-atom distance between the top 2 residues with high G-score is less than a defined distance threshold. For 90.6% (452 of 499) of the significant hits, the top G-score pair of residues are within 15 Ångstroms in 3-D space. Decreasing the distance threshold to 8 and 5 Ångstroms results in 74.7% (373) and 61.1% (305) hits whose top pairs are close in 3-D, respectively.

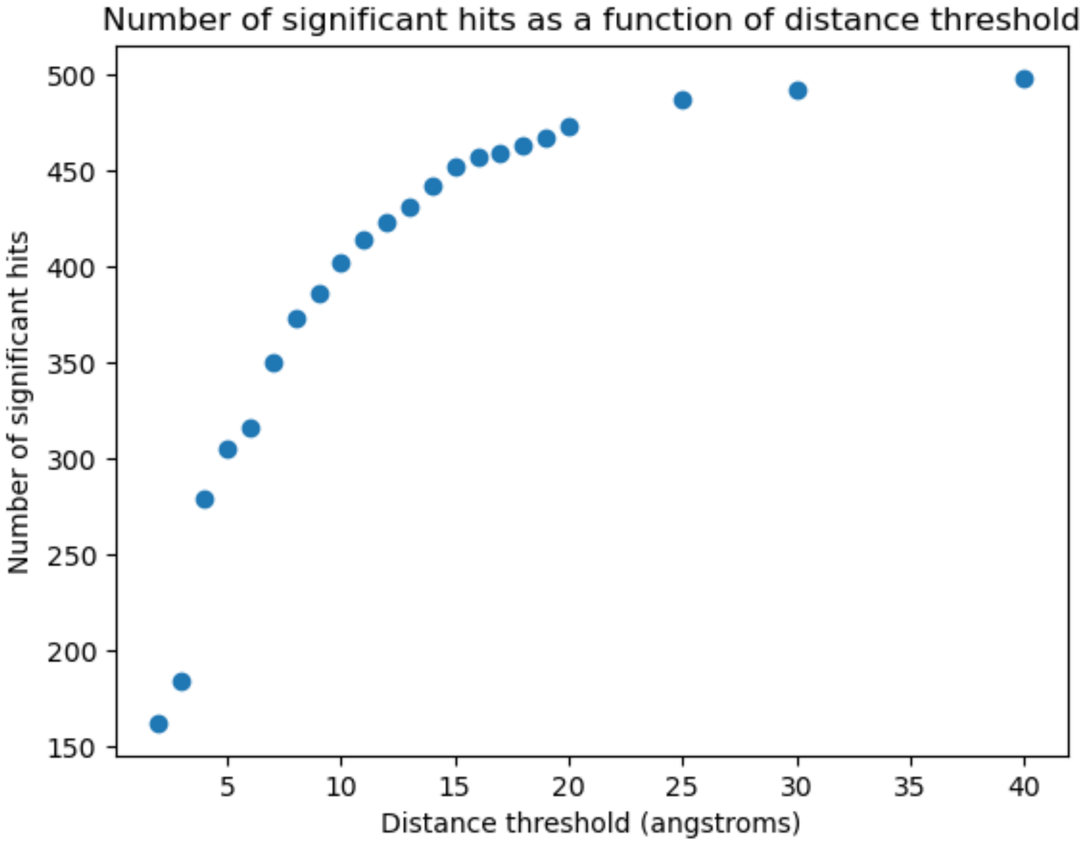

**Supplementary Figure S4: Relationship between protein length and distance between top pair of residues.** Of the 499 proteins with significant hits, we analyze whether there is a statistically significant relationship between the length of the protein (filtered for residues that pass our structure quality thresholds) and the distance between the two residues with highest G-score. Using Ordinary Least Squares regression implemented with default parameters in statsmodels v0.14.4, we find a weak negative relationship: $R^2 = 0.008$, $\beta = -3.1806$, *p*-value = 0.048, indicating that distance between top pairs does not increase with protein length, hence our method is likely capturing real signal for 3-D clustering.

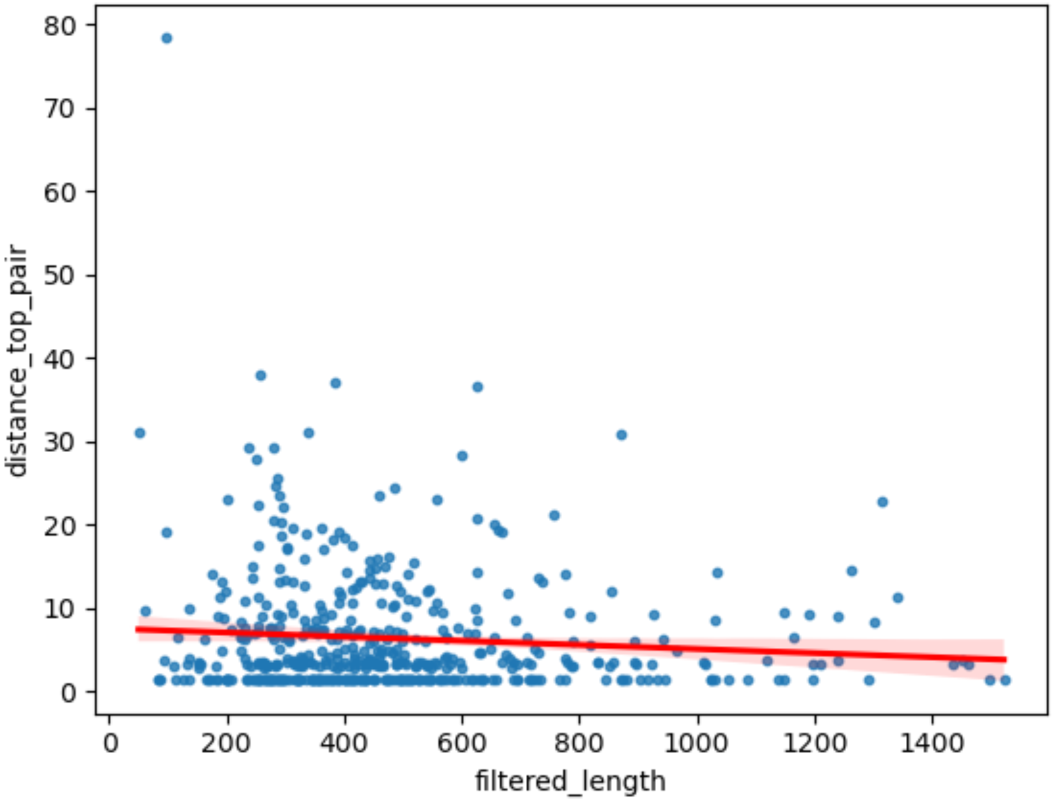

| GO id | GO term label | Uniprot identifiers | Fold enrich. | FDR | category |
|---|---|---|---|---|---|
| GO:0035635 | entry of bacterium into host cell | Q6MX51_MYCTU, GLMU_MYCTU, SAHH_MYCTU | 11.04 | 0.01 | host cell entry |
| GO:0006265 | DNA topological change | GYRA_MYCTU, GYRB_MYCTU, TOP1_MYCTU | 11.04 | 0.01 | DNA topology |
| GO:0016539 | intein-mediated protein splicing | DNAB_MYCTU, RECA_MYCTU, Y1461_MYCTU | 11.04 | 0.01 | protein maturation |
| GO:0046349 | amino sugar biosynthetic process | GLMM_MYCTU, MURA_MYCTU, GLMU_MYCTU | 11.04 | 0.01 | amino sugar |
| GO:0006047 | UDP-N-acetylglucosamine metabolic process | GLMM_MYCTU, GLMS_MYCTU, GLMU_MYCTU | 11.04 | 0.01 | amino sugar |
| GO:0062014 | negative regulation of small molecule metabolic process | PKNB_MYCTU, GARA_MYCTU, PKNA_MYCTU, PKNE_MYCTU, PKND_MYCTU | 9.20 | 0.00 | fatty acid |
| GO:0042304 | regulation of fatty acid biosynthetic process | PKNB_MYCTU, PKNA_MYCTU, PKNE_MYCTU, PKND_MYCTU | 8.83 | 0.01 | fatty acid |
| GO:0006040 | amino sugar metabolic process | GLMM_MYCTU, GLMS_MYCTU, MURA_MYCTU, GLMU_MYCTU | 8.83 | 0.01 | amino sugar |
| GO:0015990 | electron transport coupled proton transport | NUOM_MYCTU, COX1_MYCTU, NUOL_MYCTU | 8.28 | 0.04 | ETC |
| GO:0046890 | regulation of lipid biosynthetic process | PKNB_MYCTU, PKNA_MYCTU, PKNE_MYCTU, P71814_MYCTU, PKND_MYCTU | 7.88 | 0.00 | fatty acid |

**Supplementary Table 1***: Top 10 GO categories significantly enriched in the clustered protein set.* Categories with identical members and FDR (*e.g., GO:0071103 DNA conformation change and GO:0006265 DNA topological change)* have only one representative category shown. See **Supplementary Data 6** for complete table.
